# An activating mutation in AGEF-1, a putative Arf GEF, causes yolk extrusion from *C. elegans* embryos

**DOI:** 10.64898/2026.02.10.705107

**Authors:** Clare FitzPatrick, Olga Skorobogata, Ali M. Fazlollahi, Kimberley D. Gauthier, Christian E. Rocheleau

## Abstract

*C. elegans* AGEF-1, an ortholog of human ARFGEF1 and ARFGEF2, functions with ARF-1, ARF-5 and the AP-1 clathrin adaptor to regulate membrane trafficking. Similar phenotypes induced by the *agef-1(vh4[E1028K])* allele and *agef-1(RNAi)* suggested that *agef-1(vh4)* was a hypomorph. Here we report that *agef-1(vh4)* results in extrusion of yolk from the embryo. This is suppressed by RNAi of *agef-1, arf-1, arf-5* but not AP-1. Based on structure of the yeast AGEF-1 ortholog, Sec7p, the E1028K change is predicted to activate AGEF-1. We propose that Arf GTPase cycling is required to regulate trafficking with AP-1 but not with Arf effectors regulating yolk trafficking.

## Description

*C. elegans* yolk is synthesized in the hermaphrodite intestine, associates with apoB-like vitellogenins, and is then secreted into the pseudocoelom (body cavity) before being endocytosed into maturing oocytes (Grant & Hirsh, 1999; Hall et al., 1999; Kimble & Sharrock, 1983). During embryogenesis yolk granules becomes distributed amongst the dividing blastomeres. Once the primordial intestine is formed, yolk accumulates in the intestinal cells (Bossinger & Schierenberg, 1996). Little is known about how yolk is distributed during embryogenesis.

Arf GTPases are regulators of membrane trafficking that cycle between a GTP-bound “on” state and a GDP-bound “off” state regulated by guanine nucleotide exchange factors (GEFs) and GTPase activating proteins (GAPs), respectively (Jackson & Bouvet, 2014). When bound to GTP, Arfs can interact with effector proteins to regulate membrane trafficking. *C. elegans* AGEF-1 is a putative GEF orthologous to yeast Sec7p and the human ARFGEF1 and ARFGEF2 that activate class I and II Arf GTPases (Ishizaki et al., 2008; Sato et al., 2006; Skorobogata et al., 2014; Togawa et al., 1999). We previously reported that a missense allele, *agef-1(vh4)*, that results in a Glutamic acid to Lysine change in the HDS2 domain (Figure 1A) caused mislocalization of the LET-23/Epidermal Growth Factor Receptor (EGFR) in the vulva precursor cells and caused enlargement of endosomes and lysosomes in coelomocytes (Skorobogata et al., 2014). Both phenotypes were phenocopied by *agef-1(RNAi)*, and a deletion allele was zygotic lethal, suggesting that *agef-1(vh4)* was a hypomorphic allele (Skorobogata et al., 2014; Tang et al., 2012).

**Figure 1.**
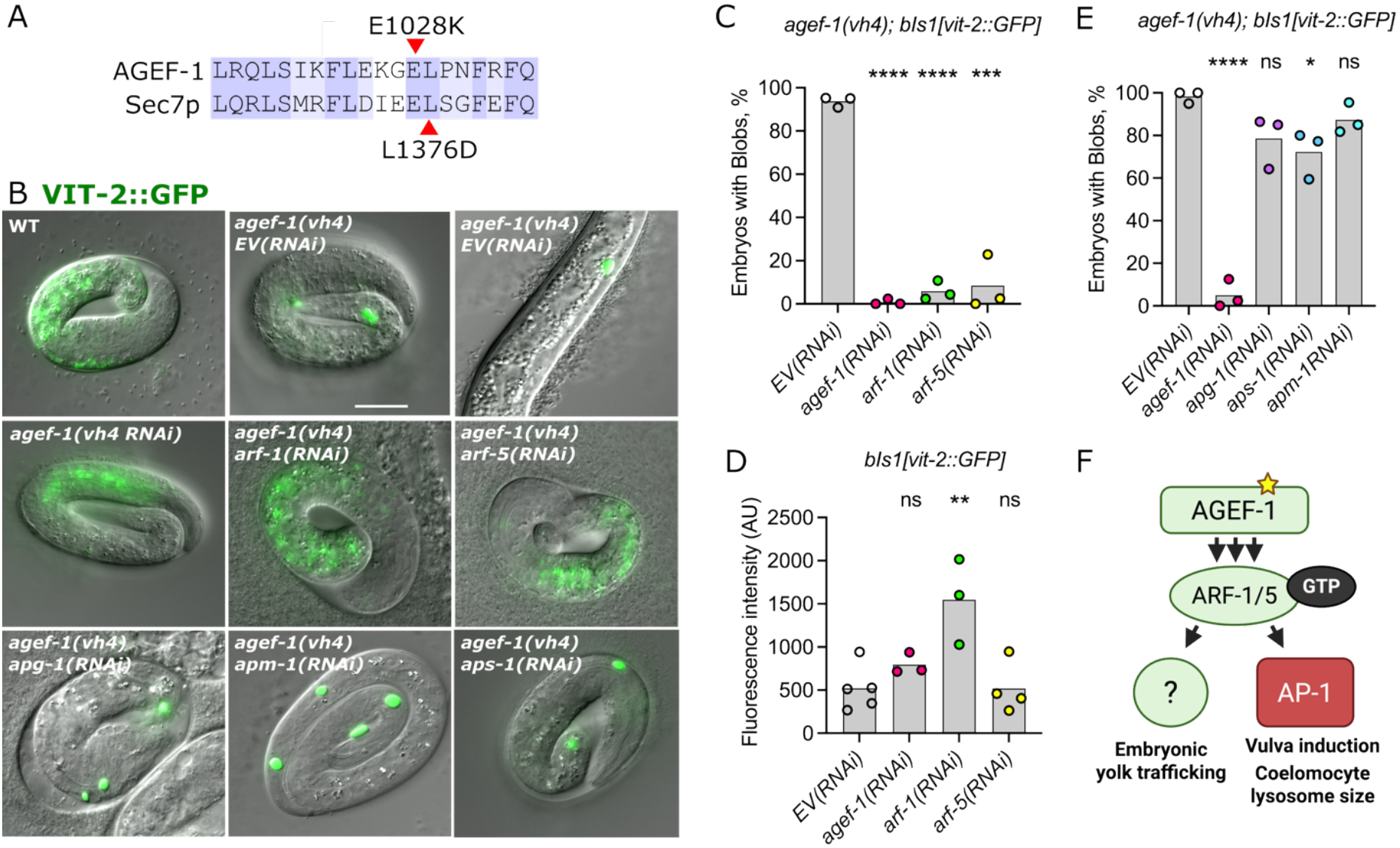
The yolk blob phenotype of *agef-1(vh4)* requires the ARF-1 and ARF-5 GTPases but not the AP-1 complex. **(A)** Alignment of amino acids from the HDS2 domains of *C. elegans* AGEF-1 and *S. cerevisiae* Sec7p. The activating L1376D mutation in Sec7p corresponds to L1029 of AGEF-1 and is adjacent to E1028 which is changed to a Lysine in the *agef-1(vh4)* mutant. Identical residues are shown in dark purple and similar residues in light purple. **(B)** Merged differential interference contrast and epifluorescence images of wild-type (WT) and *agef-1(vh4)* three-fold stage embryos and an L1 larva (upper right) expressing the VIT-2::GFP transgene, *bIs1*, and treated for RNAi targeting *agef-1, arf-1, arf-5, apg-1, apm-1, aps-1* and an empty vector (EV) control. VIT-2::GFP is localized to the intestine in wild type but is consolidated in droplets or blobs in *agef-1(vh4)* that pool between the embryo and the eggshell as well as internally as can be seen in the hatched L1 larva (upper right). RNAi targeting *agef-1, arf-1* and *arf-5* suppressed the *agef-1(vh4)* yolk blob phenotype **(C)** but did not reduce the fluorescence intensity of VIT-2::GFP in oocytes **(D)**. RNAi targeting *apg-1, apm-1* or *aps-1* did not suppress the *agef-1(vh4)* yolk blob phenotype **(E). (F)** Model that the activating mutation in AGEF-1 (star) increases the levels of active GTP-bound ARF-1 and ARF-5 (green) that can engage effectors. Since GTPase cycling is important for AP-1 mediated vesicle trafficking the net result is an inhibition of AP-1 trafficking events (red) during vulva induction and regulating lysosome size in coelomocytes. In the case of embryonic yolk trafficking Arf GTPase cycling does not appear to be required to activate an unknown effector (?; green) to inappropriately misdirect yolk. Unpaired t test was used to determine significance of percentages or fluorescence intensity. ns, not significant, * P<0.05, **P<0.01, ***P<0.001, ****P<0.0001. Number of embryos or oocytes quantified were between 102-151 (C), 29-66 (D), and 95-121 (E) per condition.

Here we report that *agef-1(vh4)* mutant embryos had a yolk trafficking phenotype. Unlike wild type, *agef-1(vh4)* embryos accumulated extraembryonic yolk in between the eggshell and the developing embryo as determined by differential interference contrast microscopy and confirmed with a yolk protein marker VIT-2::GFP (*bIs1)* (Figure 1B). In nearly all *agef-1(vh4)* embryos VIT-2::GFP was found in pools or blobs rather than in the intestine. While many yolk blobs were clearly floating between the embryo and the eggshell, some appeared to accumulate internally. Analysis of newly hatched larvae confirmed that 53% (n=41) had small yolk blobs that failed to extrude from the embryo. However, it was not clear if this phenotype was caused by loss of *agef-1* or a background mutation as we did not observe a yolk blob phenotype by *agef-1(RNAi)* (0% yolk blobs across 3 replicates, *n=*72-84 embryos/replicate).

Recent structural analysis of the yeast ortholog of AGEF-1, Sec7p, revealed that the HDS2 domain interacts with the SEC7 GEF domain, blocking the interaction with Arf1 (Brownfield et al., 2024). Consistent with this being an autoinhibitory interaction, an engineered L1376D mutation in the HDS2 domain greatly increased Sec7p GEF activity toward Arf1 *in vitro*. This Leucine is conserved in AGEF-1 and is adjacent to the Glutamic Acid that in mutated in *agef-1(vh4)* suggesting that this allele could also be an activating allele (Figure 1A). To test if *agef-1(vh4)* is indeed an activating allele, we performed *agef-1* RNAi on the *agef-1(vh4)* mutant and found that it potently suppressed the yolk blob phenotype while control empty vector (EV) RNAi had no effect (Figure 1B, C). Thus, *agef-1(vh4)* is likely a hypermorphic allele and the yolk blobs may be a result of increased Arf GTPase activity.

AGEF-1 functions with ARF-1 and ARF-5 to antagonize LET-23/EGFR localization and signaling during vulva development (Skorobogata et al., 2014). We found that RNAi of either *arf-1* or *arf-5* strongly suppressed the *agef-1(vh4)* yolk blob phenotype suggesting that both are required (Figure 1B, C). To ensure that this suppression is not caused by decreased yolk uptake into maturing oocytes we measured fluorescence intensity of VIT-2::GFP in oocytes treated with RNAi targeting *agef-1, arf-1* or *arf-5*. We found no decrease in VIT-2::GFP fluorescence intensity from these RNAi knockdowns and in fact *arf-1(RNAi)* significantly increased the levels of VIT-2::GFP (Figure 1D). Thus, ARF-1 and ARF-5 activity are required for the yolk extrusion phenotype of *agef-1(vh4)*.

The *agef-1(vh4)* mutant behaved as an activating allele in the context of yolk trafficking, but acted as a loss of function mutation during LET-23/EGFR-mediated vulva development and in regulation of lysosome size in coelomocytes (Skorobogata et al., 2014). In mammalian cells Arf1 activation is required for AP-1 recruitment (Stamnes & Rothman, 1993), but GTP hydrolysis appears to be required for AP-1 uncoating and subsequent vesicle trafficking (Meyer et al., 2005; Tanigawa et al., 1993; Zhu et al., 1998). We hypothesized that the AP-1 complex would not be required for the *agef-1(vh4)* yolk blob phenotype. We found that RNAi targeting AP-1 complex components *apg-1, apm-1* or *aps-1* did not suppress the *agef-1(vh4)* yolk blob phenotype despite causing a potent dead egg phenotype (Figure 1B, E). Therefore, the aberrant yolk trafficking seen in *agef-1(vh4)* happens independently of the AP-1 clathrin adaptor complex.

Here we demonstrated that *agef-1(vh4)* is likely a hypermorphic allele and that aberrant AGEF-1 activity results in mislocalization of yolk outside of the embryo. A similar yolk trafficking phenotype has been reported for loss *alfa-1/C9orf72* and *smcr-8/SMCR8*, where ALFA-1 and SMCR-8 were found to regulate the endolysosomal trafficking (Corrionero & Horvitz, 2018). Intriguingly, structural and biochemical data suggests that C9orf72 and SMCR8 function together as an Arf GAP (Su et al., 2021; Su et al., 2020). Further analysis will be required to determine if ALFA-1 and SMCR-8 function in a common pathway with AGEF-1 to regulate yolk trafficking.

We demonstrated a strong requirement for ARF-1 and ARF-5 downstream of AGEF-1(E1028K) induced yolk trafficking phenotypes. The strong phenotypes suggest that they may not function redundantly but possibly in a linear pathway or as a heterodimer as ARF dimerization has previously been reported (Beck et al., 2008). While we have not identified the ARF-1/5 effector, our data suggests that unlike AP-1, it will not require Arf cycling to promote trafficking. Human ARFGEF1 variants are associated with developmental delay with and without epilepsy (Takata et al., 2019; Thomas et al., 2021; Xu et al., 2022). While most ARFGEF1 mutants introduce premature stop codons and frameshifts some are missense mutations. Notably the I1180R mutation in the HDS2 domain could be an activating allele as the corresponding residue in yeast makes contact with the Sec7 GEF domain in the autoinhibited state (Brownfield et al., 2024). If so, we would expect the corresponding mutant in *agef-1* would cause a yolk trafficking phenotype.

## Methods

Wormbase (https://wormbase.org) was an invaluable resource to the planning and execution of this work (Sternberg et al., 2024). All strains were maintained at 20°C on Nematode Growth Medium (NGM) and fed HB101 *E. coli* as a food source, as previously described (Brenner, 1974; Stiernagle, 2006). All strains were derived from the wild type N2 strain. RNAi by feeding was performed as previously reported (Kamath et al., 2003). All experiments were performed a minimum of 3 times. All images were collected on an Axio Imager A1 using the Axiocam 305 with Axio Vision software (Zeiss). Embryos were mounted on 2% agarose pads in water and imaged using glass coverslips between 0.16 to 0.19 mm (Fisher). For yolk blob scoring, a minimum of one VIT-2::GFP positive blob was the threshold for the presence of a yolk blob. A percent of blobs present was taken after each round of RNAi. For total VIT-2::GFP amounts, images were captured as previously described and mean fluorescence intensity was measured using the selection tool brush on Fiji, and subtracting background (https://imagej.net/software/fiji/). Statistics and graphical analysis were performed using Graph Pad Prism 10. Sequence alignment was performed using EMBOSS Water Pairwise Sequence Alignment (PSA) (https://www.ebi.ac.uk/jdispatcher/psa/emboss_water?format=clustal). Visualization and final alignment was performed using Jalview version 2.11.4.1 (https://www.jalview.org/).

## Reagents

| Strain | Genotype | Source |
| --- | --- | --- |
| QR14 | *bIs1[Pvit-2::VIT-2::GFP; rol-6(su1006)] X* | (Chotard et al., 2010); Derived from DH1033 (Grant & Hirsh, 1999) |
| QR201 | *agef-1(vh4) I; bIs1[Pvit-2::VIT-2::GFP; rol-6(su1006)] X* | (Skorobogata et al., 2014) |

| RNAi clone | Gene | Source |
| --- | --- | --- |
| L4440 | Empty vector control | (Timmons & Fire, 1998) |
| I-2M02 | *apm-1* | (Fraser et al., 2000) |
| I-6L22 | *agef-1* | (Fraser et al., 2000) |
| I-7A02 | *apg-1* | (Fraser et al., 2000) |
| III-3A13 | *arf-1* | (Kamath et al., 2003) |
| IV-4E13 | *arf-5* | (Kamath et al., 2003) |
| V-4F01 | *aps-1* | (Kamath et al., 2003) |

## Acknowledgements

We would like to thank Jung Hwa Seo for technical assistance. Anna Corrioniro (previously of the Horvitz lab at MIT) and Chris Fromme (Cornell University) for helpful discussions. Richard Roy and Jeremy Van Raamsdonk (McGill University) for sharing reagents. The *agef-1(vh4)* allele and *bIs1* transgenic line are available at the Caenorhabditis Genetics Center (CGC), which is funded by NIH Office of Research Infrastructure Programs (P40 OD010440).

## Funding

This work was funded by a Canadian Institutes of Health Research (CIHR) Project Grant PJT-191910 to CER. CF was supported by a Canada Graduate Scholarship; OS was supported by a FRQS studentship and KDG was supported by studentships from FRQS and NSERC.

## Notes

### Competing Interest Statement

The authors have declared no competing interest.

